# Paleolithic DNA from the Caucasus reveals core of West Eurasian ancestry

**DOI:** 10.1101/423079

**Authors:** Iosif Lazaridis, Anna Belfer-Cohen, Swapan Mallick, Nick Patterson, Olivia Cheronet, Nadin Rohland, Guy Bar-Oz, Ofer Bar-Yosef, Nino Jakeli, Eliso Kvavadze, David Lordkipanidze, Zinovi Matzkevich, Tengiz Meshveliani, Brendan J. Culleton, Douglas J. Kennett, Ron Pinhasi, David Reich

## Abstract

The earliest ancient DNA data of modern humans from Europe dates to ∼40 thousand years ago^1-4^, but that from the Caucasus and the Near East to only ∼14 thousand years ago^5,6^, from populations who lived long after the Last Glacial Maximum (LGM) ∼26.5-19 thousand years ago^7^. To address this imbalance and to better understand the relationship of Europeans and Near Easterners, we report genome-wide data from two ∼26 thousand year old individuals from Dzudzuana Cave in Georgia in the Caucasus from around the beginning of the LGM. Surprisingly, the Dzudzuana population was more closely related to early agriculturalists from western Anatolia ∼8 thousand years ago^8^ than to the hunter-gatherers of the Caucasus from the same region of western Georgia of ∼13-10 thousand years ago^5^. Most of the Dzudzuana population’s ancestry was deeply related to the post-glacial western European hunter-gatherers of the ‘Villabruna cluster’^3^, but it also had ancestry from a lineage that had separated from the great majority of non-African populations before they separated from each other, proving that such ‘Basal Eurasians’^6,9^ were present in West Eurasia twice as early as previously recorded^5,6^. We document major population turnover in the Near East after the time of Dzudzuana, showing that the highly differentiated Holocene populations of the region^6^ were formed by ‘Ancient North Eurasian’^3,9,10^ admixture into the Caucasus and Iran and North African^11,12^ admixture into the Natufians of the Levant. We finally show that the Dzudzuana population contributed the majority of the ancestry of post-Ice Age people in the Near East, North Africa, and even parts of Europe, thereby becoming the largest single contributor of ancestry of all present-day West Eurasians.

---

Ancient DNA has revealed more about the deep history of Europe than of any other continent, with dozens of Paleolithic samples reported to date^1-5^ (Fig. 1a). Genetic analyses show that the first populations related to present-day West Eurasians arrived in Europe at least ∼36 thousand years ago (kya)^2^. A new group of populations (Věstonice cluster), associated with the archaeologically defined Gravettian entity, appeared in the genetic record of Europe by ∼30kya, while another group, associated with the archaeologically defined Magdalenian culture, appeared in Europe by ∼20kya (El Mirón cluster) ^3^. By ∼14kya a third group, the Villabruna cluster, appeared throughout mainland Europe, coinciding with the Bølling-Allerød warming period^3^. Members of this cluster, which has also been called western European hunter-gatherers (WHG), were found across Europe during Late Upper Paleolithic-to-Mesolithic times, and were the main pre-agricultural Europeans prior to the Neolithic ∼8kya^9^.

**Figure 1.**
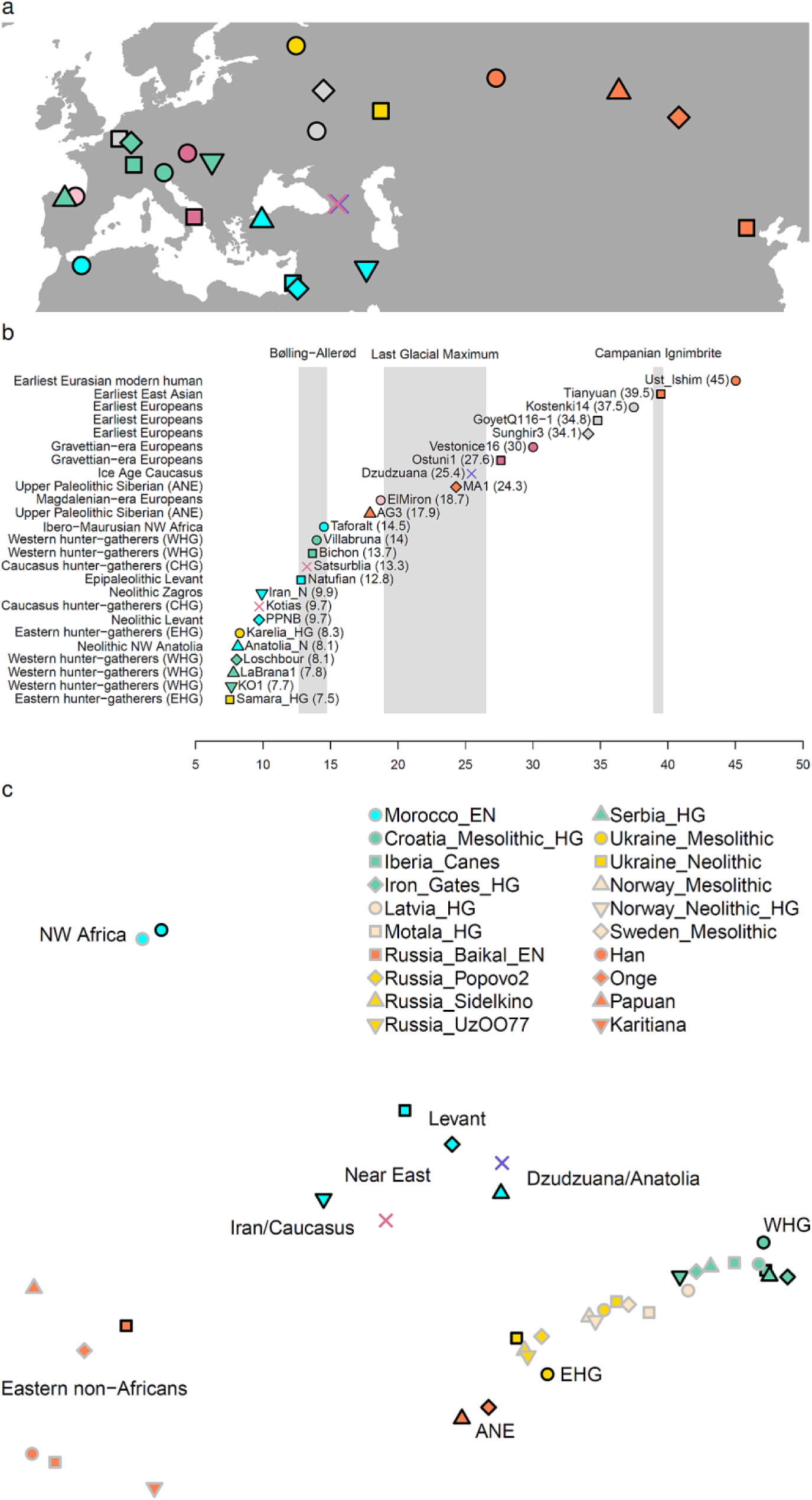
Ancient West Eurasian population structure. (a) Geographical distribution of key ancient West Eurasian populations. (b) Temporal distribution of key ancient West Eurasian populations (approximate date in ky BP). (c) PCA of key ancient West Eurasians, including additional populations (shown with grey shells), in the space of outgroup *f*_4_-statistics (Methods).

In contrast to this detailed knowledge about Europe during the Paleolithic, no Ice Age DNA has been published from the Near East (including the Caucasus) whose post-glacial and Holocene-era populations <15kya were highly differentiated from both those of Europe and also from each other^5,6,8,13,14^. To address this deficit, we analyzed teeth from two individuals recovered from Dzudzuana Cave^15^, Southern Caucasus, (Fig. 1a; Supplementary Information section 1) from an archaeological layer previously dated to ∼27-24kya and whose age determination was confirmed by a series of 8 new dates (Extended Data Figure 1; Supplementary Information section 1), thus allowing us to probe a population on the cusp of the beginning of the LGM. Of the two individuals, one yielded data at a total of 185,744 targeted single nucleotide polymorphisms (SNPs) on chromosomes 1-22 (I2949, Dzu2, Upper Area square H16b), while the other yielded 90,916 SNPs (I2963, Dzu3, Lower Area square 18b). Both individuals had mitochondrial DNA sequences (U6 and N) that are consistent with deriving from lineages that are rare in the Caucasus or Europe today. The two individuals were genetically similar to each other, consistent with belonging to the same population (Extended Data Fig. 2) and we thus analyze them jointly.

The Dzudzuana samples represent the earliest ancient modern human DNA outside of Europe, Siberia, and China (Fig. 1b). In the local context, they help us answer the question of the relationship of Ice Age populations of the region to their post-glacial successors^5^: was there discontinuity in the Caucasus as in Europe? A broader issue that we wished to address is the changing relationship between human populations from Europe and the Near East, as the Caucasus is situated at the border between them. The Villabruna cluster has been modeled as contributing to both the ∼30kya Věstonice and ∼20kya El Mirón-cluster populations^3^, suggesting that it must have existed somewhere in relatively unmixed form long before the oldest genetic data we have from it at ∼14kya^3,5^. However, it is unlikely that the Villabruna cluster sojourned in mainland Europe, as members of the cluster have been attested there only by ∼14kya, marking an increased affinity of these European populations of the time to Near Eastern ones^3^. Was there migration at the time from mainland Europe to the Near East or vice versa, or, indeed from a geographically intermediate Ice Age refugium in southeast Europe, Anatolia, or the circum-Pontic (Black Sea) region that might explain the affinity of post-glacial Levantine and Anatolian populations to those of Europe^6^? It is also unknown how the affinity between early populations in the eastern European-Caucasus-Iran zone^6^ first arose. Eastern European hunter-gatherers (EHG)^16^ ∼8kya can be modeled as a mixture of peoples of WHG and Upper Paleolithic Siberians first known ∼24kya^10^ (also known as ‘Ancient North Eurasians’ (ANE)). Caucasus hunter-gatherers (CHG)—sampled in Georgia in Satsurblia and Kotias Klde caves <50km from Dzudzuana^5^—were genetically intermediate between EHG and the first agriculturalists of Iran sampled from the Zagros mountains (Iran_N; ∼10kya)^6,13^.

We first estimated *F*_ST_, a measure of population genetic differentiation, to assess the genetic relationships between ancient West Eurasian populations (Extended Data Table 1; Methods). Post-glacial Near Easterners and North Africans (PGNE) (CHG, Natufians, Taforalt^11^ Ibero-Maurusians from North Africa, and early Neolithic farmers from Anatolia^8^, Iran^6^, the Levant^6^, and the Maghreb^17^) are strongly differentiated from all European and Siberian hunter-gatherers (ESHG) (*F*_ST_ = 0.078-0.267). By contrast, Dzudzuana is genetically closer to both contemporaneous Gravettians from Europe (0.051±0.012) and also to the much later Neolithic Anatolian farmers (0.039±0.005) who are genetically closest to them according to this measure. Genetic drift inflates *F*_ST_ over time, so the affinity to the Gravettians may partly be due to the great age of these samples. However, age cannot explain the affinity to much later Neolithic Anatolians of ∼8kya, a population closer to Dzudzuana than any other PGNE (0.052-0.195).

Outgroup *f*_3_-statistics^10^ show that Dzudzuana clusters with Near Eastern populations primarily from Anatolia and secondarily from the Levant, but not with the geographically proximate CHG (Extended Data Fig. 3). A genetic relationship between Dzudzuana and Neolithic Anatolians is also shown by principal components analysis (PCA) in the space of ‘outgroup *f*_4_-statistics’^16^ of the form *f*_4_(*Test, O*_1_; *O*_2_, *O*_3_) where (*O*_1_; *O*_2_, *O*_3_) is a triple of outgroups (Fig. 1c; Methods); performing PCA on the space defined by these statistics has the advantage of not being affected by genetic drift peculiar to the *Test* populations. It also allows us to visualize genetic relationships between ancient populations alone, without projecting onto the variation of present-day people. European hunter-gatherers in our analysis form a cline with Villabruna/WHG samples on one end and ANE on the other. None of the PGNE populations other than the Neolithic Anatolians cluster with the Ice Age Caucasus population from Dzudzuana. As reported previously, present-day West Eurasians are much more homogeneous than ancient ones, reflecting extensive post-Neolithic admixture^6^. However, they continue to be differentially related to ancient local populations in Europe and the Near East (Extended Data Fig. 4).

To better understand the relationship of Dzudzuana to other ancient West Eurasian populations, we performed symmetry testing using *f*-statistics^18^ (Extended Data Fig. 5). These analyses show that ESHG share more alleles with Dzudzuana than with PGNE populations, except Neolithic Anatolians who form a clade with Dzudzuana to the exclusion of ESHG (Extended Data Fig. 5a). Thus, our results prove that the European affinity of Neolithic Anatolians^6^ does not necessarily reflect any admixture into the Near East from Europe, as an Anatolian Neolithic-like population already existed in parts of the Near East by ∼26kya. Furthermore, Dzudzuana shares more alleles with Villabruna-cluster groups than with other ESHG (Extended Data Fig. 5b), suggesting that this European affinity was specifically related to the Villabruna cluster, and indicating that the Villabruna affinity of PGNE populations from Anatolia and the Levant is not the result of a migration into the Near East from Europe. Rather, ancestry deeply related to the Villabruna cluster was present not only in Gravettian and Magdalenian-era Europeans^3^ but also in the populations of the Caucasus, by ∼26kya. Neolithic Anatolians, while forming a clade with Dzudzuana with respect to ESHG (Extended Data Fig. 5a), share more alleles with all other PGNE (Extended Data Fig. 5d), suggesting that PGNE share at least partially common descent to the exclusion of the much older samples from Dzudzuana.

All known ancient Near Eastern populations prior to this work were inferred to harbor ‘Basal Eurasian’ ancestry^9^, a branch that diverged from all other non-Africans (including ESHG and present-day East Asians and Oceanians) before they split from each other. The CHG, geographically intermediate between Europe and the Near East resembled Near Eastern populations in the possession of Basal Eurasian ancestry^5^. The Dzudzuana population was not identical to the WHG, as it shared fewer alleles with both an early Upper Paleolithic Siberian (Ust’Ishim^19^) and an early Upper Paleolithic East Asian (Tianyuan^20^) (Extended Data Fig. 5c), thus, it too—like the PGNE populations—had Basal Eurasian ancestry^6,9^. The detection of this type of ancestry, twice as early as previously documented^5,6^ and at the northern edge of the Near East, lends weight to the hypothesis that it represents a deep Near Eastern lineage rather than a recent arrival from Africa^6^.

We used qpGraph^18^ to build an admixture graph model of the relationship between ESHG and Dzudzuana, also including the earliest PGNE populations from North Africa (Taforalt) and the Epipaleolithic Levant (Natufians) (Fig. 2). While potentially oversimplifying the history of these populations by considering only discrete binary admixture events as opposed to continuous gene flow, the model is useful for its insights into possible evolutionary relationships between populations and for representing the minimum complexity that these relationships had. According to this model, a common population contributed ancestry to Gravettians (represented by Vestonice16) and to a “Common West Eurasian” population that contributed all the ancestry of Villabruna and most of the ancestry of Dzudzuana which also had 28.4±4.2% Basal Eurasian ancestry^21^ (Supplementary Information section 2).

**Figure 2.**
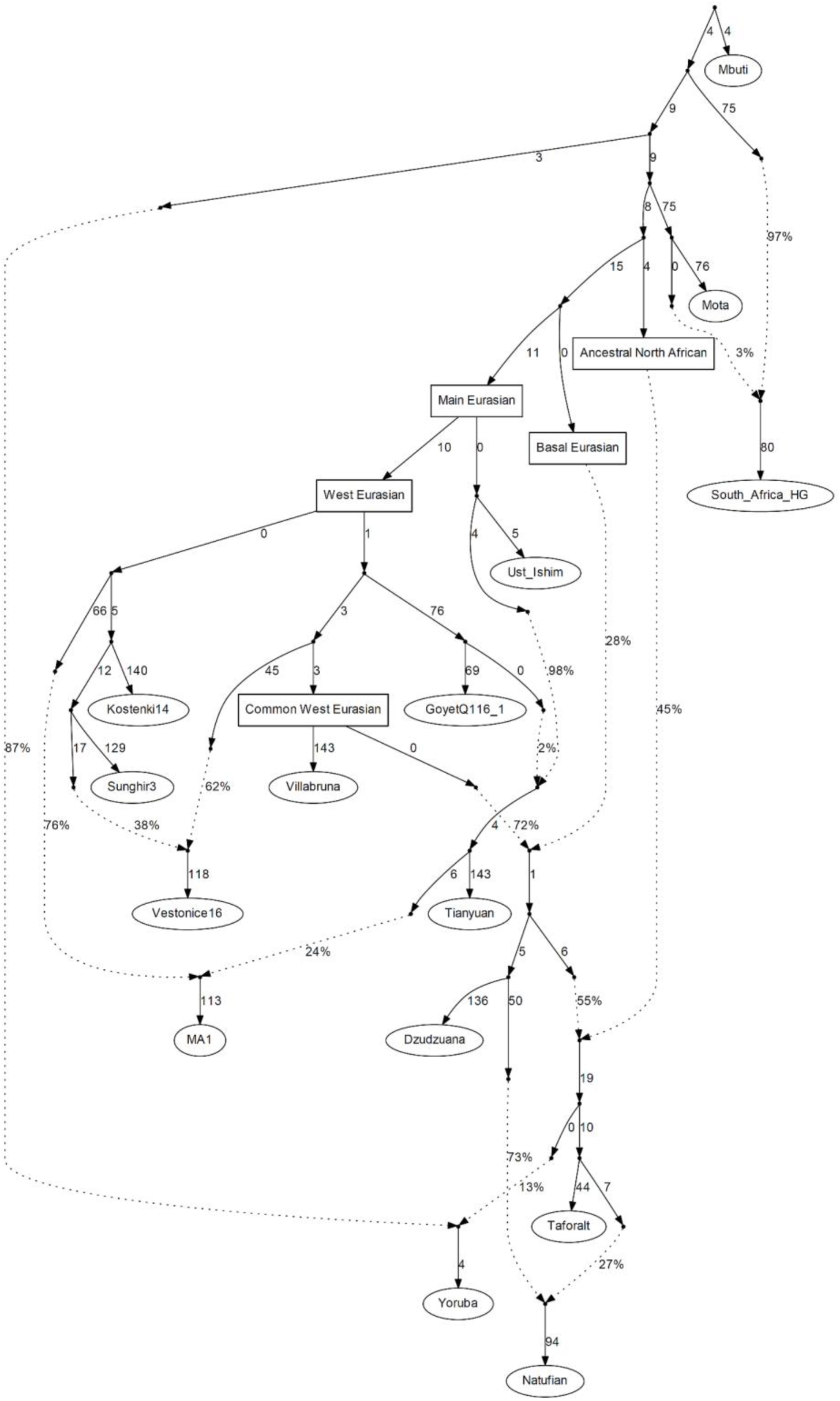
An admixture graph model of Paleolithic West Eurasians. An automatically generated admixture graph models fits 14 populations (worst Z-score of the difference between estimated and fitted *f*-statistics is 2.7) or 15 populations (also including South_Africa_HG, worst Z-score is 3.5). This is a simplified model assuming binary admixture events and is not a unique solution (Supplementary Information section 2). Sampled populations are shown with ovals and select labeled internal nodes with rectangles.

Our co-modeling of Epipaleolithic Natufians and Ibero-Maurusians from Taforalt confirms that the Taforalt population was mixed^11^, but instead of specifying gene flow from the ancestors of Natufians into the ancestors of Taforalt as originally reported, we infer gene flow in the reverse direction (into Natufians). The Neolithic population from Morocco, closely related to Taforalt^17^ is also consistent with being descended from the source of this gene flow, and appears to have no admixture from the Levantine Neolithic (Supplementary Information section 3). If our model is correct, Epipaleolithic Natufians trace part of their ancestry to North Africa, consistent with morphological and archaeological studies that indicate a spread of morphological features^22^ and artifacts from North Africa into the Near East. Such a scenario would also explain the presence of Y-chromosome haplogroup E in the Natufians and Levantine farmers^6^, a common link between the Levant and Africa. Moreover, our model predicts that West Africans (represented by Yoruba) had 12.5±1.1% ancestry from a Taforalt-related group rather than Taforalt having ancestry from an unknown Sub-Saharan African source^11^; this may have mediated the limited Neanderthal admixture present in West Africans^23^. An advantage of our model is that it allows for a local North African component in the ancestry of Taforalt, rather than deriving them exclusively from Levantine and Sub-Saharan sources.

We also used the qpWave/qpAdm framework^16^ to model ancient populations without strong phylogenetic assumptions (Supplementary Information section 3; Table 1). This analysis shows that we cannot reject the hypothesis that Dzudzuana and the much later Neolithic Anatolians form a clade with respect to ESHG (P=0.286), consistent with the latter being a population largely descended from Dzudzuana-like pre-Neolithic populations whose geographical extent spanned both Anatolia and the Caucasus. Dzudzuana itself can be modeled as a 2-way mixture of Villabruna-related ancestry and a Basal Eurasian lineage. Western PGNE populations, including Neolithic Anatolians, pre-pottery Neolithic farmers from the Levant (PPNB), Natufians, and Taforalt, can all be modeled as a mixture of Dzudzuana and additional ‘Deep’ ancestry that may represent an even earlier split than the Basal Eurasians. Considering 2-way mixtures, we can model Karelia_HG as deriving 34±2.8% of its ancestry from a Villabruna-related source, with the remainder mainly from ANE represented by the AfontovaGora3 (AG3) sample from Lake Baikal^3^ ∼17kya. Finally, we can model CHG and samples from Neolithic Iran (Iran_N) as deriving their ancestry largely (∼58-64% using qpAdm and ∼45-62% using qpGraph) from a Dzudzuana-like population, but with ancestry from both ‘Deep’ and ANE sources, thus proving that ANE ancestry had reached Western Eurasia long before the Bronze Age Eurasian steppe migrations that carried further westward into mainland Europe.

**Table 1:**
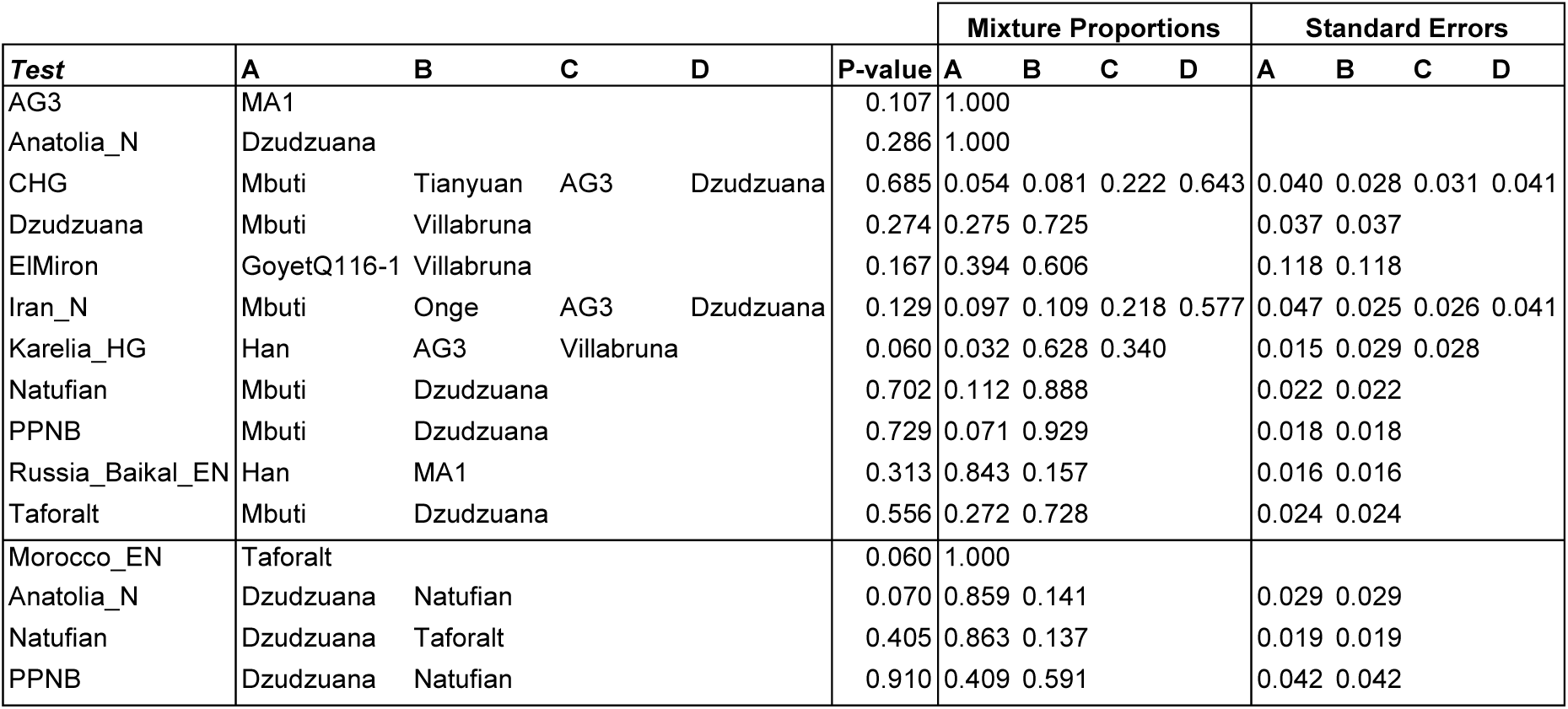
Mixture proportions of ancient populations. The model with the fewest sources for each population that fits the data is shown. Multiple models may fit some populations; we show here the one with the highest P-value; others are shown in Supplementary Information section 3. The P-value is (for *N* sources) for rank *N*-1 of (Test, Sources) with respect to a set of diverse outgroups (Supplementary Information section 3). Neolithic Near Eastern populations (four bottom rows) can also be fit as mixtures of Dzudzuana and Taforalt via Natufians.

| Test | A | B | C | D | P-value | Mixture Proportions |  |  |  | Standard Errors |  |  |  |
| --- | --- | --- | --- | --- | --- | --- | --- | --- | --- | --- | --- | --- | --- |
|  |  |  |  |  |  | A | B | C | D | A | B | C | D |
| AG3 | MA1 |  |  |  | 0.107 | 1.000 |  |  |  |  |  |  |  |
| Anatolia_N | Dzudzuana |  |  |  | 0.286 | 1.000 |  |  |  |  |  |  |  |
| CHG | Mbuti | Tianyuan | AG3 | Dzudzuana | 0.685 | 0.054 | 0.081 | 0.222 | 0.643 | 0.040 | 0.028 | 0.031 | 0.041 |
| Dzudzuana | Mbuti | Villabruna |  |  | 0.274 | 0.275 | 0.725 |  |  | 0.037 | 0.037 |  |  |
| ElMiron | GoyetQ116-1 | Villabruna |  |  | 0.167 | 0.394 | 0.606 |  |  | 0.118 | 0.118 |  |  |
| Iran_N | Mbuti | Onge | AG3 | Dzudzuana | 0.129 | 0.097 | 0.109 | 0.218 | 0.577 | 0.047 | 0.025 | 0.026 | 0.041 |
| Karelia_HG | Han | AG3 | Villabruna |  | 0.060 | 0.032 | 0.628 | 0.340 |  | 0.015 | 0.029 | 0.028 |  |
| Natufian | Mbuti | Dzudzuana |  |  | 0.702 | 0.112 | 0.888 |  |  | 0.022 | 0.022 |  |  |
| PPNB | Mbuti | Dzudzuana |  |  | 0.729 | 0.071 | 0.929 |  |  | 0.018 | 0.018 |  |  |
| Russia_Baikal_EN | Han | MA1 |  |  | 0.313 | 0.843 | 0.157 |  |  | 0.016 | 0.016 |  |  |
| Taforalt | Mbuti | Dzudzuana |  |  | 0.556 | 0.272 | 0.728 |  |  | 0.024 | 0.024 |  |  |
| Morocco_EN | Taforalt |  |  |  | 0.060 | 1.000 |  |  |  |  |  |  |  |
| Anatolia_N | Dzudzuana | Natufian |  |  | 0.070 | 0.859 | 0.141 |  |  | 0.029 | 0.029 |  |  |
| Natufian | Dzudzuana | Taforalt |  |  | 0.405 | 0.863 | 0.137 |  |  | 0.019 | 0.019 |  |  |
| PPNB | Dzudzuana | Natufian |  |  | 0.910 | 0.409 | 0.591 |  |  | 0.042 | 0.042 |  |  |

In qpAdm modeling, a deeply divergent hunter-gatherer lineage that contributed in relatively unmixed form to the much later hunter-gatherers of the Villabruna cluster is specified as contributing to earlier hunter-gatherer groups (Gravettian Vestonice16: 35.7±11.3% and Magdalenian ElMiron: 60.6±11.3%) and to populations of the Caucasus (Dzudzuana: 72.5±3.7%, virtually identical to that inferred using ADMIXTUREGRAPH). In Europe, descendants of this lineage admixed with pre-existing hunter-gatherers related to Sunghir3 from Russia^4^ for the Gravettians and GoyetQ116-1 from Belgium^3^ for the Magdalenians, while in the Near East it did so with Basal Eurasians. Later Europeans prior to the arrival of agriculture were the product of re-settlement of this lineage after ∼15kya in mainland Europe, while in eastern Europe they admixed with Siberian hunter-gatherers forming the WHG-ANE cline of ancestry (Fig. 1c). In the Near East, the Dzudzuana-related population admixed with North African-related ancestry in the Levant and with Siberian hunter-gatherer and eastern non-African-related ancestry in Iran and the Caucasus. Thus, the highly differentiated populations at the dawn of the Neolithic^6^ were primarily descended from Villabruna Cluster and Dzudzuana-related ancestors, with varying degrees of additional input related to both North Africa and Ancient North/East Eurasia whose proximate sources may be clarified by future sampling of geographically and temporally intermediate populations.

The ancestry of present-day Europeans has been traced to the proximate sources of Mesolithic hunter-gatherers, Early European/Anatolian farmers, and steppe pastoralists^16^, but the ancestry of Near Eastern and North African populations has not been investigated due to lack of appropriate ancient sources. We present a unified analysis of diverse European, Near Eastern, North African populations in terms of the deepest known sources of ancestry (Fig. 3), which suggests that Dzudzuana-related ancestry makes up ∼46-88% of the ancestry of all these populations, with Dzudzuana-related ancestry more strongly found in southern populations across West Eurasia (Fig. 3; Extended Data Fig. 6). Dzudzuana-like ancestry must have spread across West Eurasia with Neolithic migrations out of the Near East, but it had not been previously completely absent from Europe as several hunter-gatherer populations in southeastern Europe, eastern Europe, and Scandinavia can only be modeled with some such ancestry (Extended Data Fig. 6; Supplementary Information section 4). Both Europeans and Near Easterners also share in AG3-related ancestry of up to ∼30% in eastern Europe down to ∼0% in parts of North Africa. Europeans are differentiated by an excess of up to ∼20% Villabruna-related ancestry relative to non-European populations and also by a relative lack of extra ‘Deep’ ancestry compared to the Near East and North Africa, a type of ancestry that may only partially be explained by the Basal Eurasian ancestry of ancient West Eurasian populations and must also trace to Africa (Extended Data Fig. 7). ‘Deep’ ancestry, including Basal Eurasian ancestry, is associated with reduced Neandertal ancestry (Supplementary Information section 5, Extended Data Fig. 8), confirming that Neandertal ancestry in West Eurasia^6^ has been diluted by admixture.

**Figure 3:**
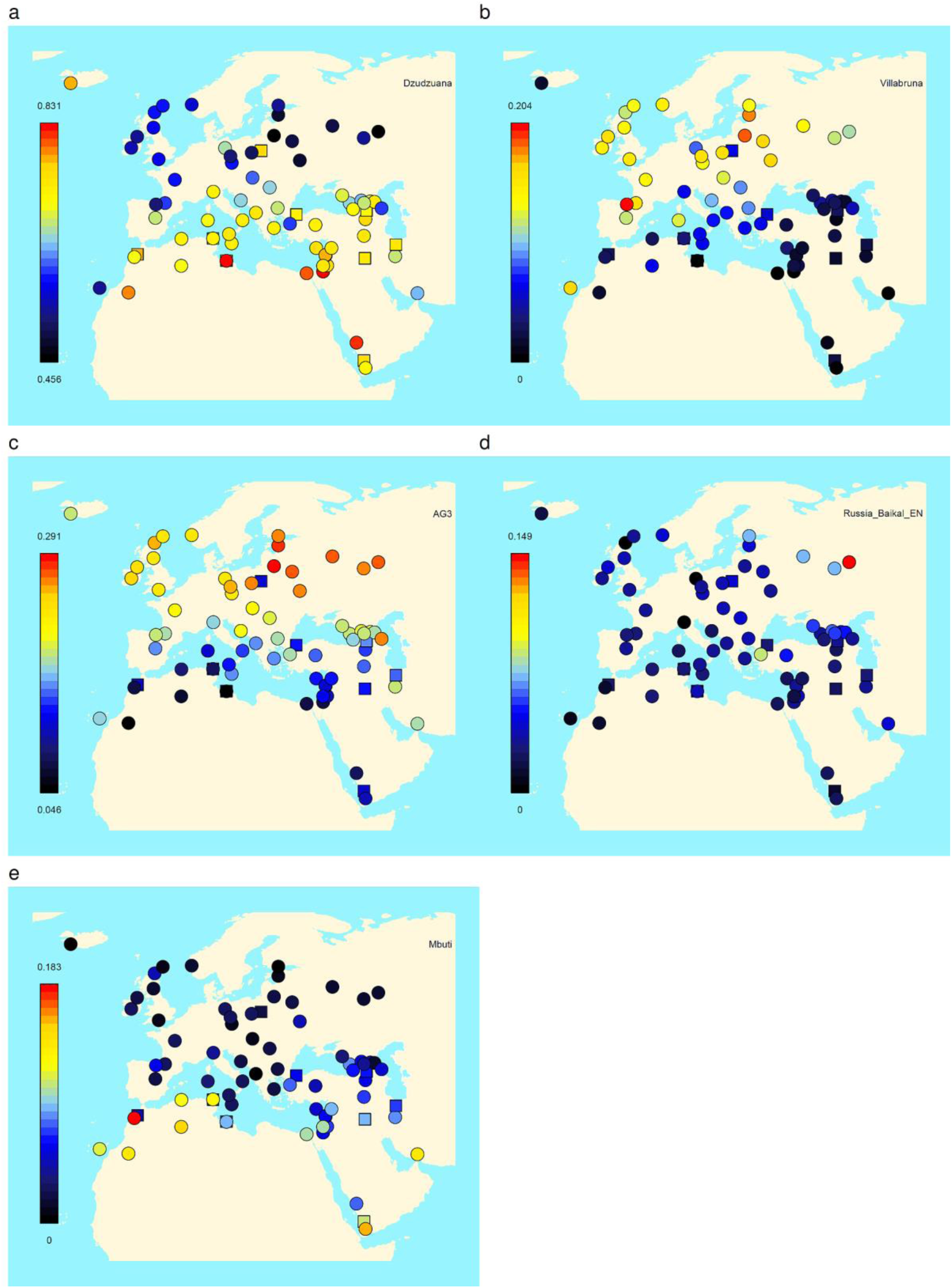
Modeling present-day West Eurasians and North Africans. Mixture proportions can be found in Extended Data Fig. 6 and Supplementary Information section 4. Ancestry with Mbuti as a source reflects all ‘Deep’ ancestry from lineages that split off prior to the 45kya Ust’Ishim. Jewish populations are shown with squares to visually distinguish them from geographically proximate populations.

Future studies must investigate when Dzudzuana-like populations first formed: does the Basal Eurasian component represent the earliest Near Eastern population stratum or a later accretion? What were the mechanisms and proximate sources of the Siberian-and North African-related ancestry that affected West Eurasia? We caution that the inference of Dzudzuana-related ancestry as the core component of ancient and present-day West Eurasia does not constitute proof for migrations specifically from the Caucasus: given that this is the only ancient DNA data from this time period and broad region, the geographical and temporal extent of this population and its relatives remains unknown. Both in its past (formed by admixture with Basal Eurasians), and in its future (admixing with populations from Africa, Europe, and Siberia in post-glacial, Neolithic, and later periods), Dzudzuana stands in the middle of an ongoing process of admixture of diverse elements from which West Eurasians, in all their diversity, eventually formed.

---

**Extended Data Figure 1:**
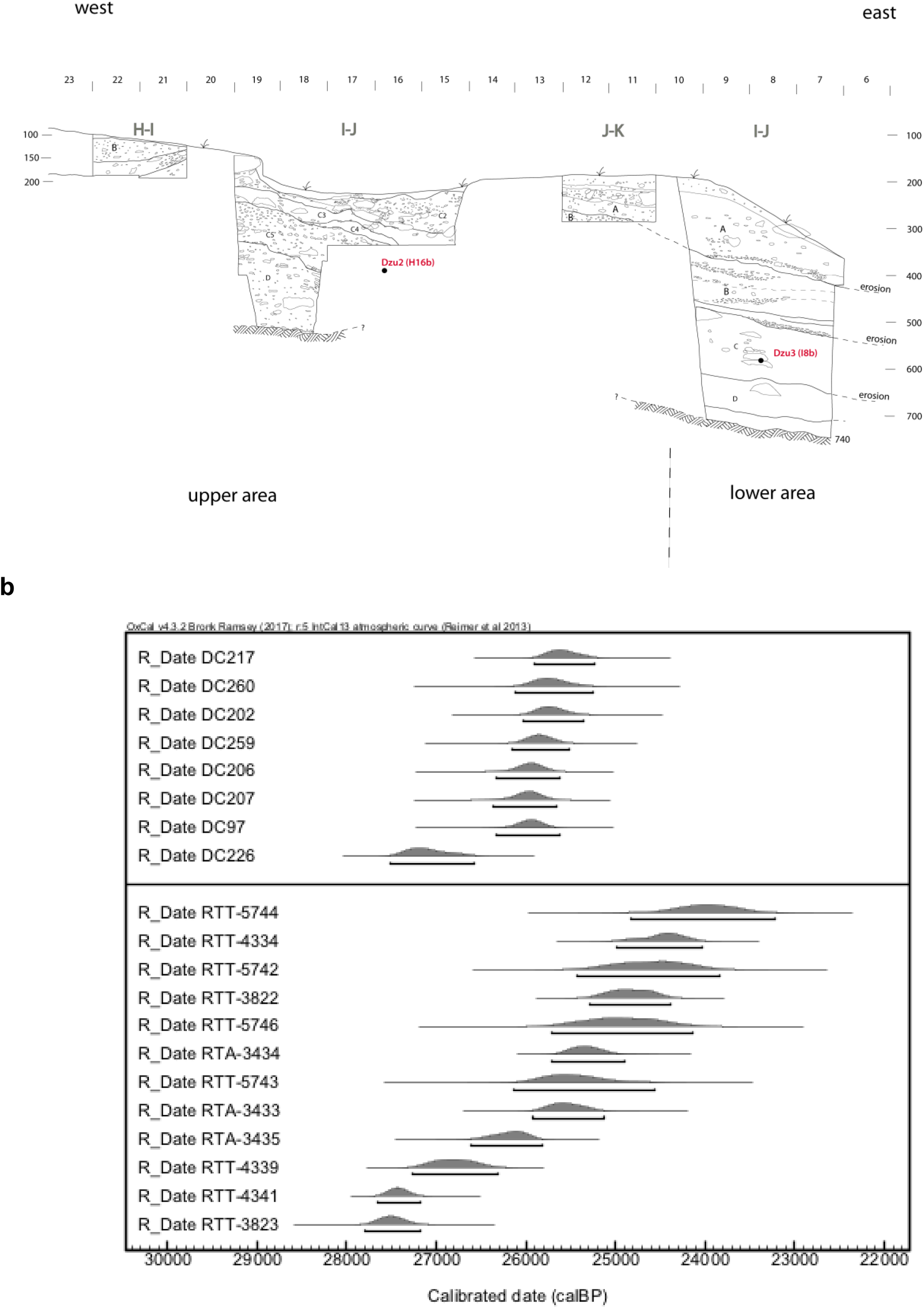
Dzudzuana Cave. (a) Section drawing of Dzudzuana Cave. Location of the two recovered teeth is indicated. (b) Calibrated dates for Layer C, upper pane: new determinations (Supplementary Information section 1), lower pane: dates from ref.^24^

**Extended Data Figure 2:**
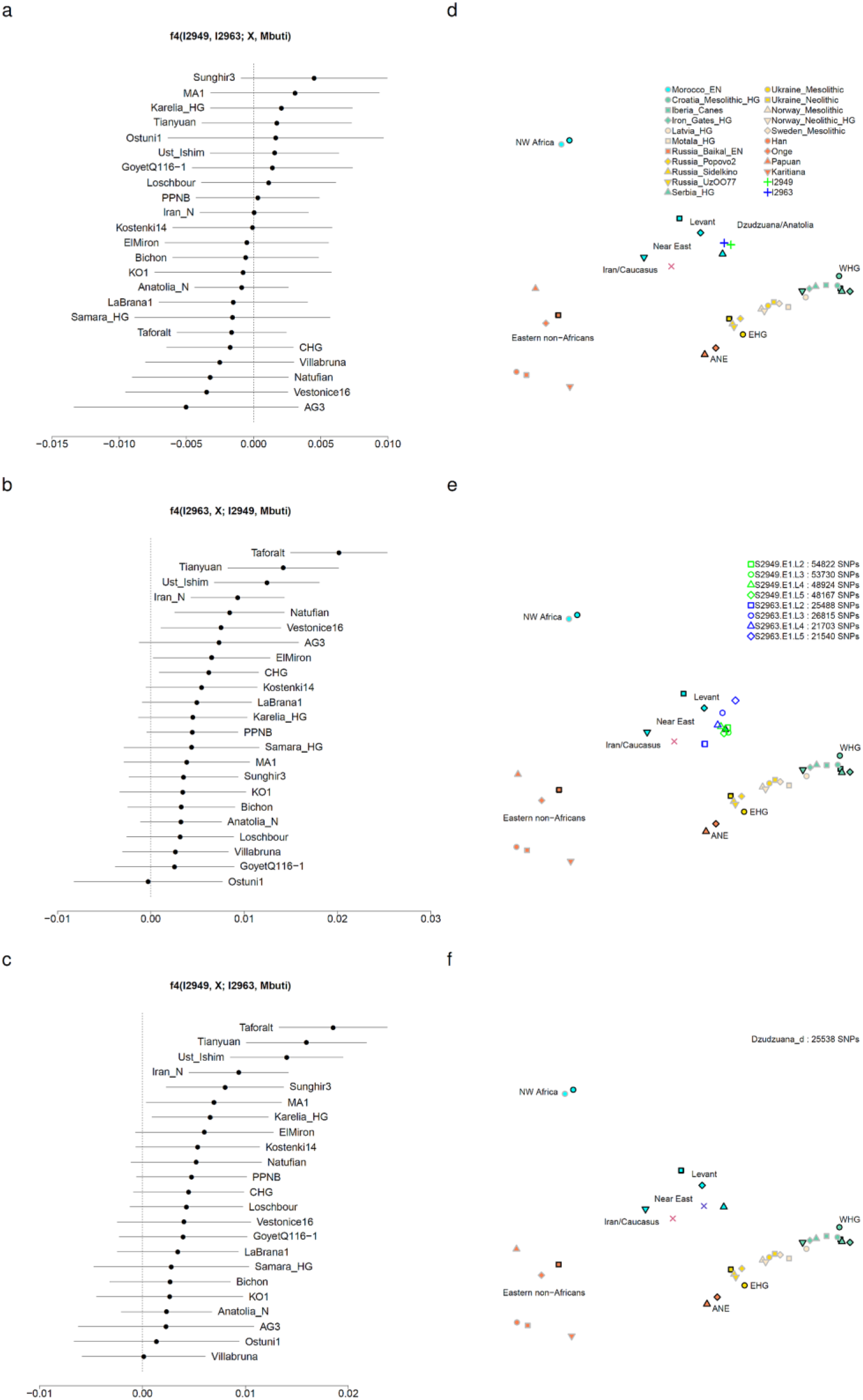
Symmetry testing and quality control. (a) The two Dzudzuana individuals are symmetrically related to others (|Z|<2.5). (b) Individual I2949 shares more alleles with I2963 than with others (Z∈[-0.1, 11.6]). (c) Individual I2963 shares more alleles with I2949 than with others (Z∈[0.1, 10.5]). (d) Principal components analysis. We repeat the analysis of Fig. 1c with the two Dzudzuana individuals shown separately. (e) Analysis of the 8 libraries separately. (f) Analysis of damage restricted sequences; point shapes and colors not shown in legend are identical to Fig. 1.

**Extended Data Figure 3:**
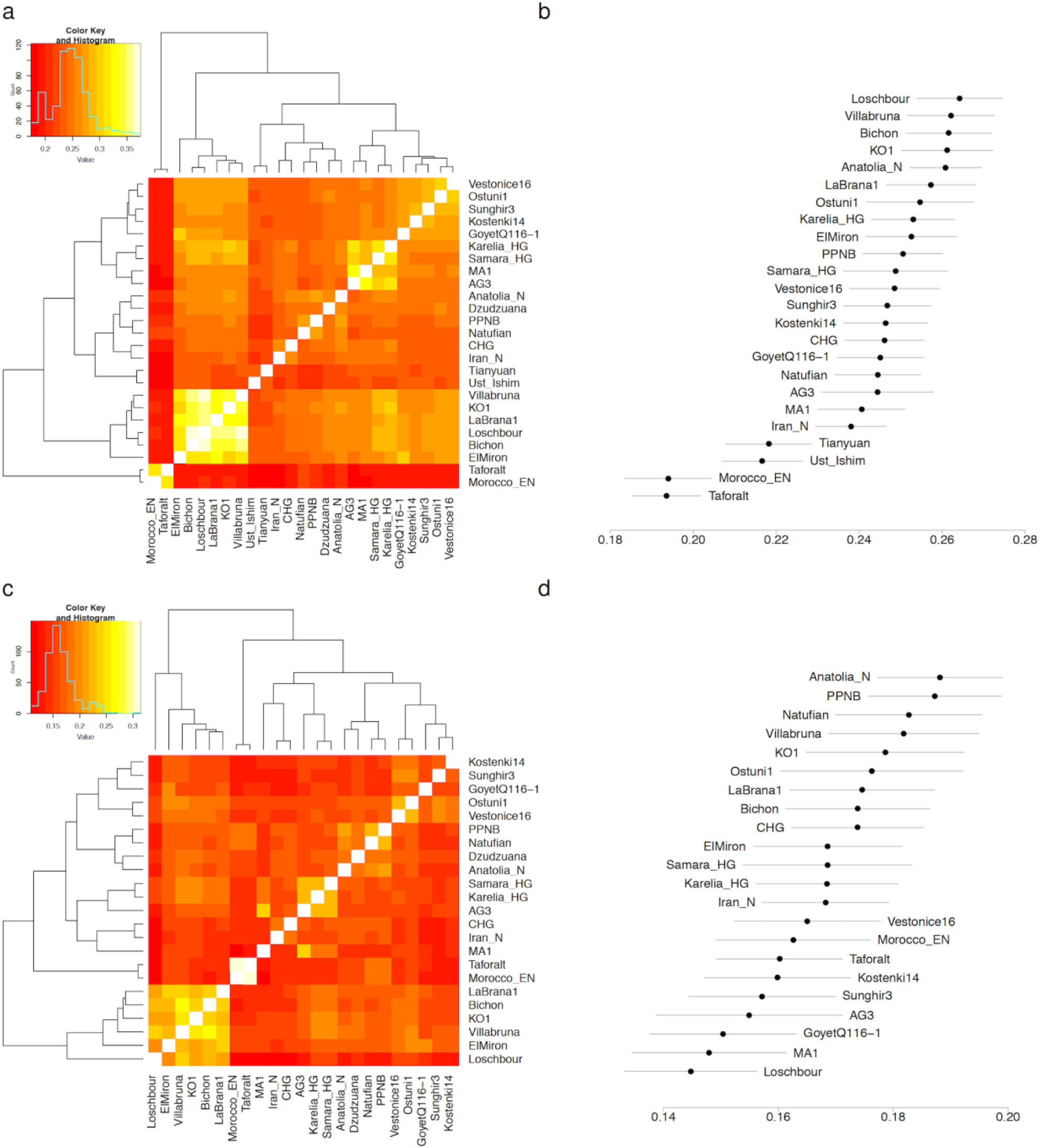
Outgroup *f*_3_-statistics of ancient West Eurasians. (a) We plot pairwise *f*_3_(Mbuti; *X, Y*) for pairs of ancient West Eurasian populations, which measure the shared genetic drift between *X* and *Y* after their separation from an African outgroup (Mbuti pygmies). (b) The value of the statistic of panel (a) for *X*=Dzudzuana with ±3 standard errors. (c) We plot *f*_3_(Ust’Ishim and Tianyuan; *X, Y*), which measures the shared genetic drift between *X* and *Y* after their separation from non-West Eurasians (made up of the Ust’Ishim and Tianyuan individuals of ∼40-45kya age). The value of the statistic of panel (c) for *X*=Dzudzuana with ±3standard errors.

**Extended Data Figure 4:**
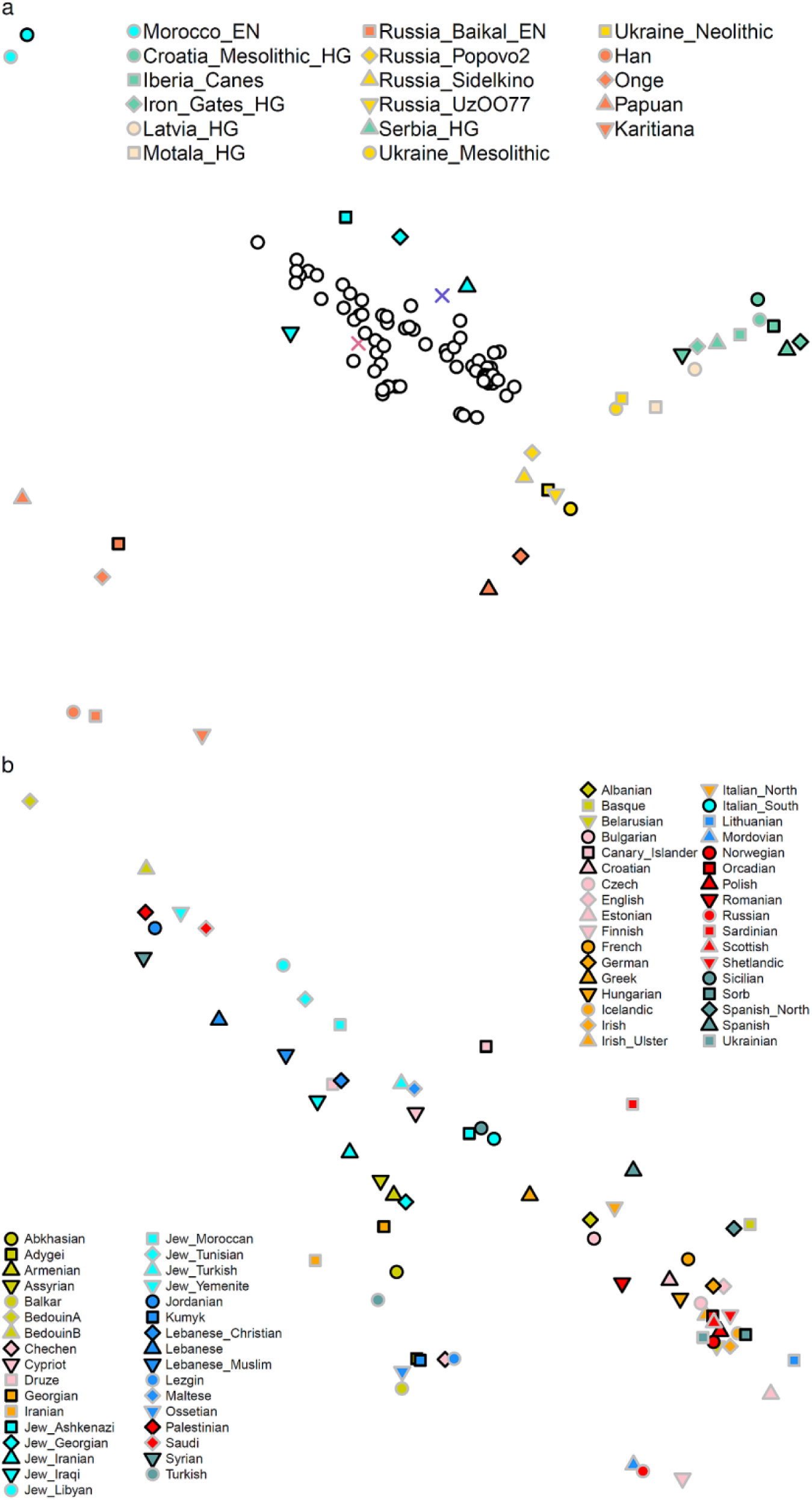
Principal components analysis of ancient and present-day West Eurasians. (a) View of all samples; present-day populations shown in white-filled circles. This corresponds to Fig. 1c, but is computed on Human Origins data. (b) View of present-day samples; the legend is split by the median PC1 value and shows a correspondence between European (bottom-right) and Near Eastern (top-left) present-day populations. Point shapes and colors not shown in legend are identical to Fig. 1.

**Extended Data Figure 5:**
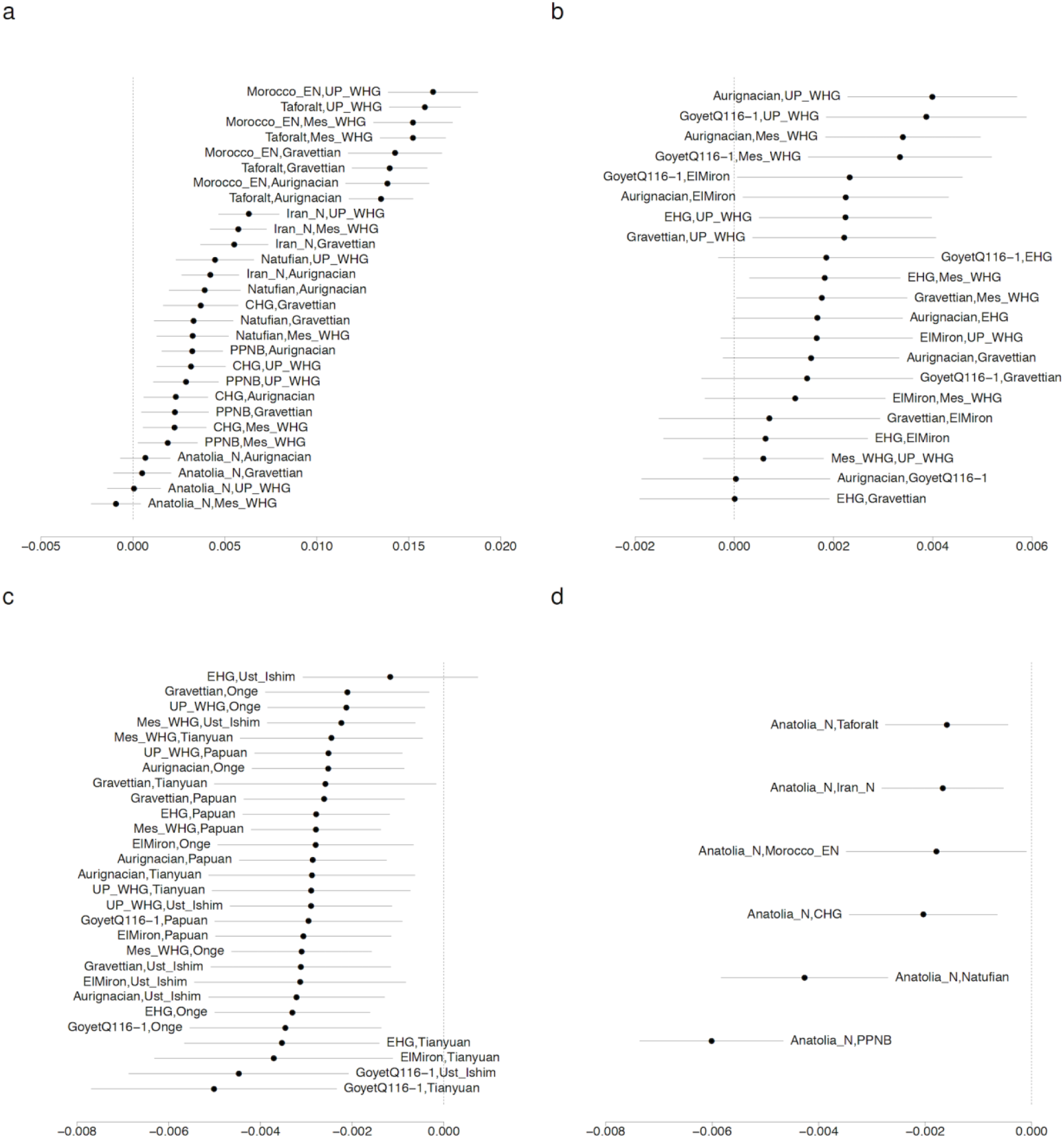
Symmetry testing. In all panels we plot the pair (*X, Y*) as a label for each statistic which is shown with ±3 standard errors (a) The statistic *f*_4_(Dzudzuana, *X, Y*; Mbuti) shows that diverse European hunter-gatherers *Y* share more alleles with Dzudzuana than with other ancient Near Eastern populations except for Neolithic Anatolians. (b) The statistic *f*_4_(Dzudzuana, Mbuti; *X, Y*) shows that Dzudzuana shares more alleles with WHG (both from the Upper Paleolithic and Mesolithic) than with other European hunter-gatherers. (c) The statistic *f*_4_(Dzudzuana, *X*; *Y*, Mbuti) shows that early Eurasians like Ust’Ishim, Tianyuan and eastern non-Africans like Onge and Papuans share more alleles with European hunter-gatherers than with Dzudzuana. (d) The statistic *f*_4_(Dzudzuana, Anatolia_N; *Y*, Mbuti) shows that Near Easterners *Y* share more alleles with Neolithic Anatolians than with Dzudzuana.

**Extended Data Figure 6:**
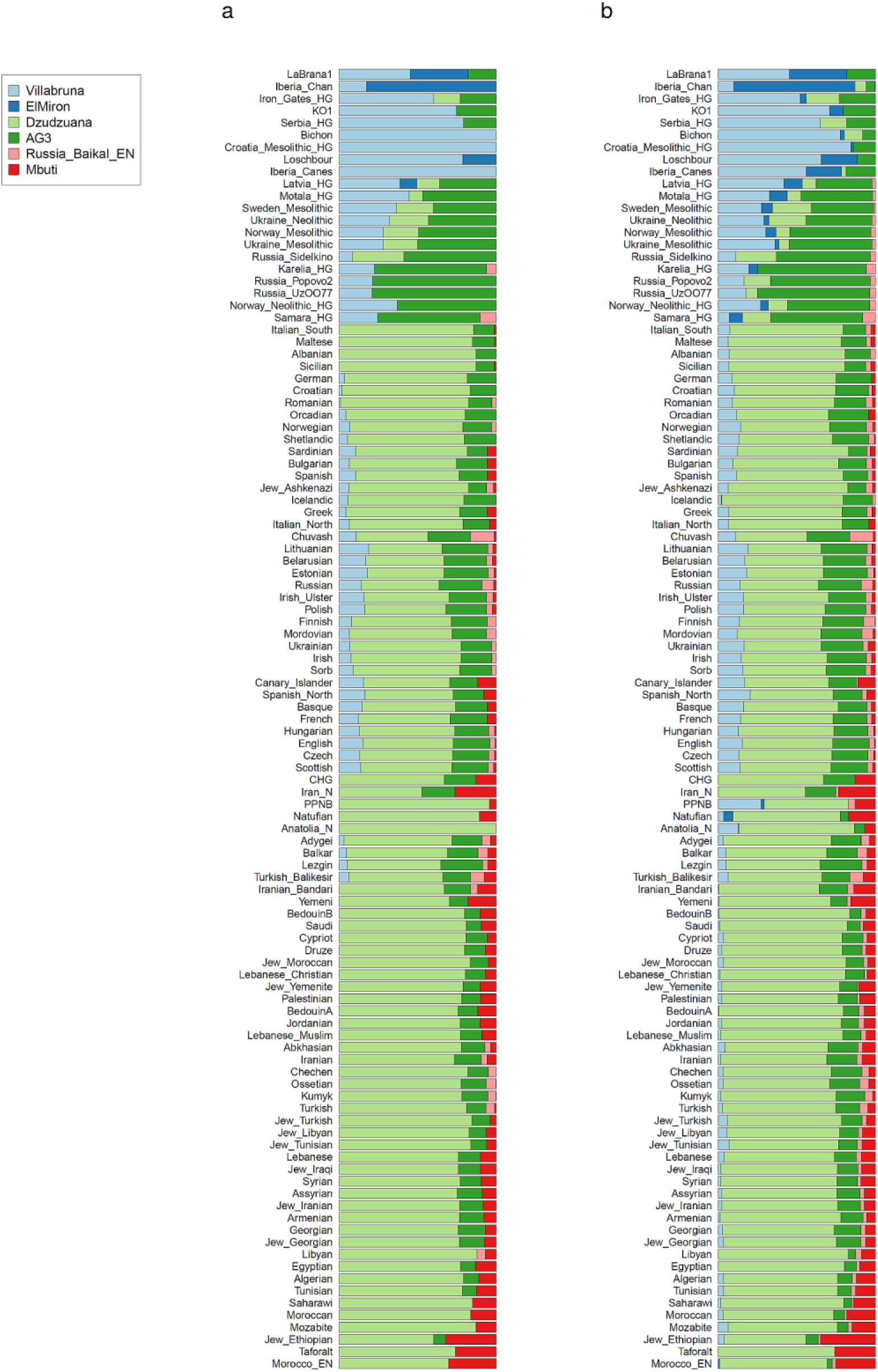
Modeling present-day and ancient West-Eurasians. Mixture proportions computed with qpAdm (Supplementary Information section 4). The proportion of ‘Mbuti’ ancestry represents the total of ‘Deep’ ancestry from lineages that split prior to the split of Ust’Ishim, Tianyuan, and West Eurasians and can include both ‘Basal Eurasian’ and other (e.g., Sub-Saharan African) ancestry. (a) ‘Conservative’ estimates. Each population cannot be modeled with fewer admixture events than shown. (b) ‘Speculative’ estimates. The highest number of sources (≤5) with admixture estimates within [0,1] are shown for each population. Some of the admixture proportions are not significantly different from 0 (Supplementary Information section 4).

**Extended Data Figure 7:**
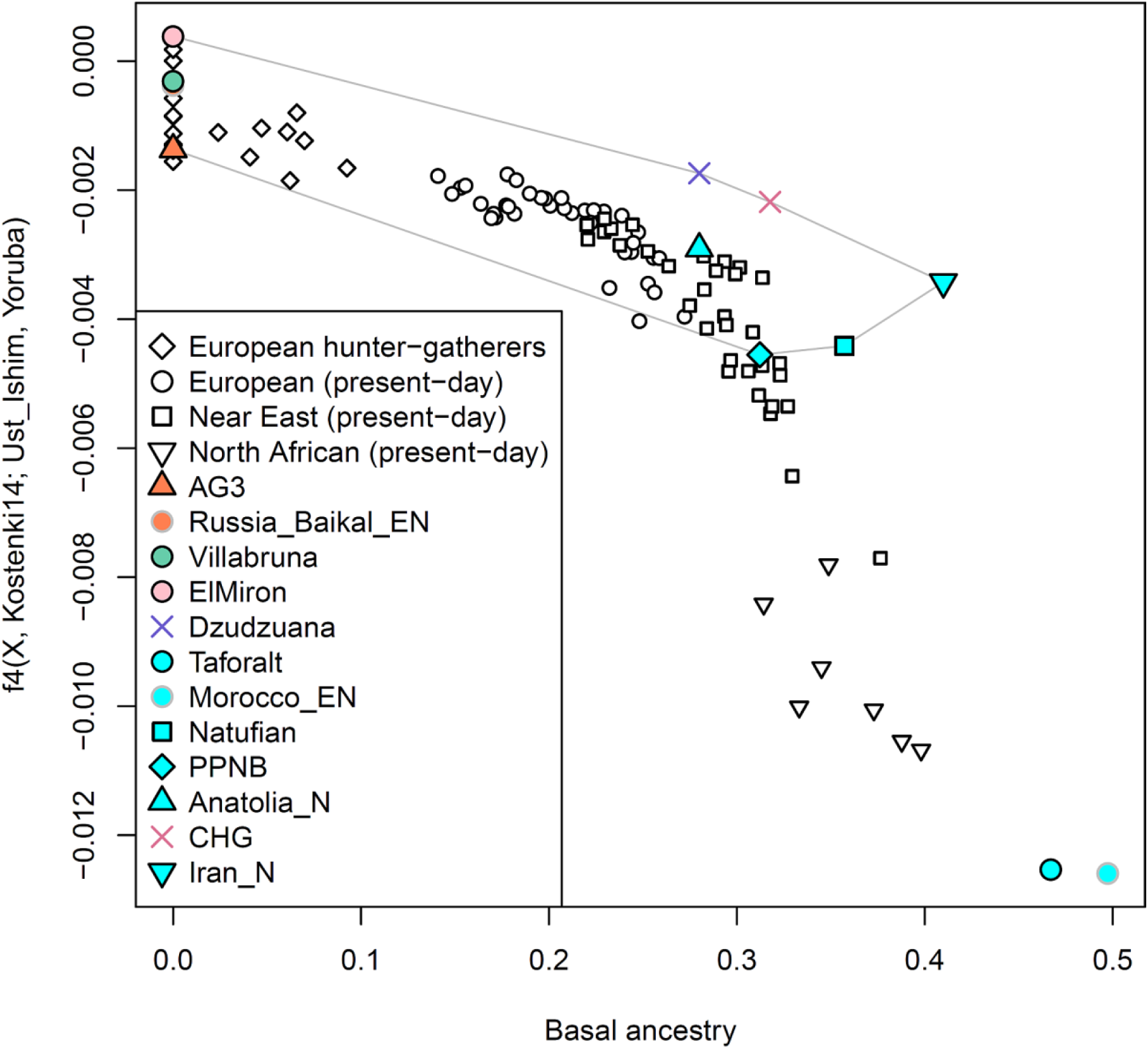
Differential relationship of Basal ancestry to Africa. Basal ancestry (conservative estimate) is negatively correlated with the statistic *f*_4_(*X,* Kostenki14, Ust’Ishim, Yoruba) which quantifies allele sharing between *X* and Ust’Ishim, consistent with this type of ancestry diluting the affinity of populations to this 45kya Siberian (earliest known modern human for which there are genomic data). For Taforalt and some populations from the Near East and North Africa this statistic is more negative, suggesting that they have North or Sub-Saharan-related ancestry that cannot be accounted for by any combination of the ancient West Eurasian sources whose convex hull is shown.

**Extended Data Figure 8:**
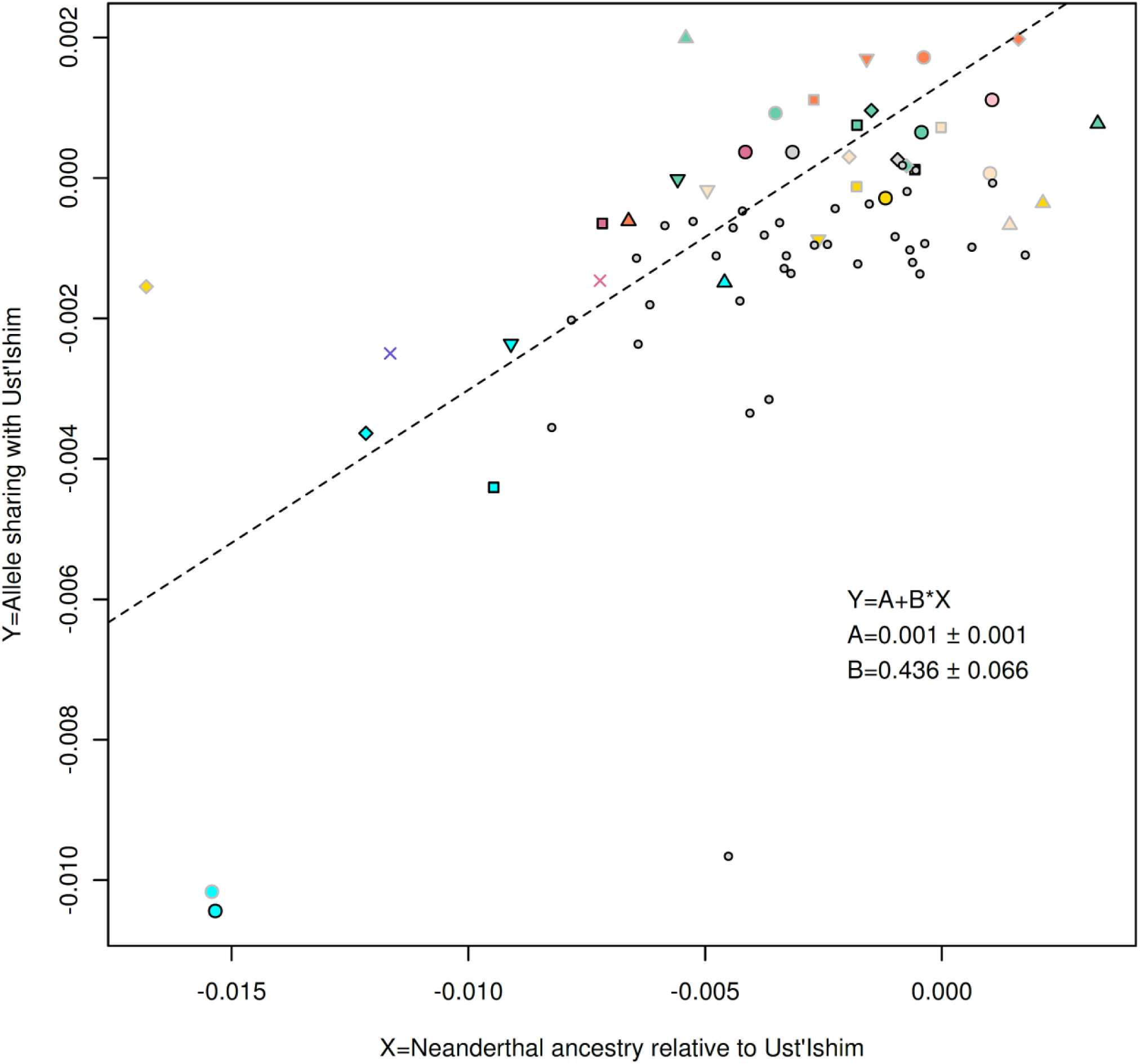
Deep ancestry of West Eurasians is associated with reduced Neandertal ancestry. Symbols used as in Fig. 1. Small grey-filled circles represent present-day West Eurasian individuals sequenced in the Simons Genome Diversity Project. The three left-most ancient samples and all the present-day populations are not used in the regression. Standard errors of the linear regression line are computed with a block jackknife (Supplementary Information section 5).

**Extended Data Table 1.** *F*_ST_ of ancient West Eurasian populations. The following groups were used as *F*_ST_ estimation requires at least two individuals per population. ANE=MA1^10^ and AG3^3^. Aurignacian-era: Kostenki14 and Sunghir3^4^. CHG: Kotias and Satsurblia^5^. EHG: Karelia_HG and Samara_HG^8,16^. Gravettian-era: Vestonice16 and Ostuni1^3^. Mes_WHG: Loschbour^9^, KO1^25^, and LaBrana1^26^. UP_WHG: Villabruna^3^ and Bichon^5^. The value of the statistic is shown below the diagonal and its standard error above it.

|  | Anatolia_N | ANE | Aurignacian | CHG | Dzudzuana | EHG | Gravettian | Iran_N | Mes_WHG | Natufian | PPNB | Taforalt | UP_WHG | Morocco_EN |
| --- | --- | --- | --- | --- | --- | --- | --- | --- | --- | --- | --- | --- | --- | --- |
| Anatolia_N |  | 0.003 | 0.002 | 0.002 | 0.005 | 0.002 | 0.002 | 0.001 | 0.002 | 0.002 | 0.001 | 0.002 | 0.003 | 0.003 |
| ANE | 0.119 |  | 0.004 | 0.004 | 0.015 | 0.004 | 0.005 | 0.003 | 0.003 | 0.005 | 0.004 | 0.003 | 0.004 | 0.007 |
| Aurignacian | 0.078 | 0.096 |  | 0.003 | 0.009 | 0.003 | 0.004 | 0.003 | 0.003 | 0.003 | 0.003 | 0.003 | 0.004 | 0.004 |
| CHG | 0.074 | 0.115 | 0.094 |  | 0.010 | 0.003 | 0.004 | 0.002 | 0.003 | 0.004 | 0.003 | 0.003 | 0.003 | 0.004 |
| Dzudzuana | 0.039 | 0.100 | 0.060 | 0.068 |  | 0.009 | 0.012 | 0.007 | 0.007 | 0.013 | 0.008 | 0.006 | 0.009 | 0.021 |
| EHG | 0.089 | 0.047 | 0.089 | 0.100 | 0.094 |  | 0.004 | 0.002 | 0.003 | 0.004 | 0.003 | 0.003 | 0.003 | 0.005 |
| Gravettian | 0.089 | 0.116 | 0.028 | 0.117 | 0.051 | 0.100 |  | 0.003 | 0.003 | 0.005 | 0.003 | 0.003 | 0.004 | 0.006 |
| Iran_N | 0.074 | 0.110 | 0.093 | 0.059 | 0.070 | 0.099 | 0.114 |  | 0.002 | 0.002 | 0.002 | 0.002 | 0.003 | 0.003 |
| Mes_WHG | 0.098 | 0.132 | 0.087 | 0.126 | 0.084 | 0.081 | 0.092 | 0.132 |  | 0.003 | 0.002 | 0.002 | 0.003 | 0.003 |
| Natufian | 0.076 | 0.164 | 0.121 | 0.131 | 0.106 | 0.142 | 0.131 | 0.126 | 0.148 |  | 0.003 | 0.003 | 0.004 | 0.007 |
| PPNB | 0.022 | 0.122 | 0.087 | 0.085 | 0.052 | 0.100 | 0.090 | 0.080 | 0.109 | 0.043 |  | 0.002 | 0.003 | 0.004 |
| Taforalt | 0.195 | 0.242 | 0.204 | 0.219 | 0.192 | 0.228 | 0.218 | 0.214 | 0.242 | 0.199 | 0.179 |  | 0.003 | 0.003 |
| UP_WHG | 0.112 | 0.149 | 0.100 | 0.139 | 0.097 | 0.096 | 0.100 | 0.144 | 0.001 | 0.160 | 0.125 | 0.252 |  | 0.004 |
| Morocco_EN | 0.206 | 0.254 | 0.219 | 0.230 | 0.195 | 0.236 | 0.234 | 0.225 | 0.252 | 0.218 | 0.190 | 0.101 | 0.267 |  |

## Methods

### Ancient DNA

In a dedicated ancient DNA clean room facility at University College Dublin, we prepared powder from the roots of teeth, and then sent these powders to Harvard Medical School where in another clean room we extracted DNA^27^,^28^ and prepared double-stranded libraries^29^,^30^ which we enriched for sequenced overlapping the mitochondrial genome and 1.24 million SNPs in the nuclear genome^1,16,31^ (Methods). We obtained usable data for two individuals after quality control and merging of data from 4 libraries for each of the two samples (treated with the enzyme UDG to reduce ancient DNA errors). The sequences from all libraries displayed characteristic damage in the terminal nucleotides consistent with partial UDG treatment.

### Contamination testing

We assessed contamination by examining heterozygosity on mitochondrial DNA using contamMix^19^ and schmutzi^32^ (Supplementary Data Table 1).

## Datasets

Our main analysis dataset included 1,233,013 SNPs, of which the 1,150,639 ones on the autosomes were analyzed. Present-day populations from the Simons Genome Diversity Panel^33^ were included in this dataset. Analyses of Supplementary Information section 4 that included present-day populations^6,9,13,18^ genotyped on the Affymetrix Human Origins array were performed on a dataset of 597,573 SNPs (593,124 on the autosomes).

### Estimation of *F*_ST_

We estimated *F*_ST_ in smartpca^34^ using the default parameters and inbreed: YES^35^ and fstonly: YES.

### *f*-statistics

All *f*-statistics were computed in ADMIXTOOLS^18^ using the programs qpDstat (with parameter f4mode: YES) and qp3Pop and default parameters.

### Principal component analysis on outgroup *f*_4_-statistics

We computed for *n* populations *Test*_1_, *Test*_2_,…,*Test*_n_ and *n*_o_ outgroups, a matrix of *n*×*d* dimensionality where 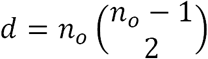 of outgroup *f*_4_-statistics^16^ of the form *f*_4_(*Test*_i_, *O*_1_; *O*_2_, *O*_3_). The outgroup set was: Vindija^36^, Altai^23^, Denisova^37^, Mbuti^33^, South_Africa_HG^38^, Mota^39^, Yoruba^33^, Han^33^, Onge^33^, Papuan^33^, Karitiana^33^, Ust_Ishim^19^, Tianyuan^20^, MA1^10^, Kostenki14^3^, GoyetQ116-1^3^, Sunghir3^4^, Vestonice16^3^, Ostuni1^3^, ElMiron^3^, Dzudzuana. PCA on the matrix was performed using the R package ppca^40^,^41^ which allows for missing data, thus allowing us to also plot populations that are included in the outgroup set. ppca was run with parameters 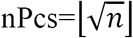 (the number of principal components used to fill in missing values), and seed=123. Results are shown in Fig. 1c and Extended Data Fig. 4.

### Admixture graph fitting

We used ADMIXTUREGRAPH program qpGraph with parameters outpop: NULL, blgsize: 0.05, forcezmode: YES, lsqmode: YES, diag: .0001, bigiter: 6, hires: YES, allsnps: YES, lambdascale: 1. We used the hash value computed by qpreroot to avoid evaluating equivalent graphs. We fit graphs in an automated way described in Supplementary Information section 2.

### qpWave/qpAdm analyses

We used qpWave^16,35^,^42^ and qpAdm^16^ to test for the number of streams of ancestry from a set of ‘right’ populations to a set of ‘left’ ones which includes the population whose history of admixture we study, and to also estimate mixture proportions (Supplementary Information section 3).

## Acknowledgments

We thank M. Lipson, I. Mathieson, N. Nakatsuka, and I. Olalde for discussions or comments on a draft of this paper. We thank N. Adamski, M. Ferry, M. Michel, J. Oppenheimer, K. Stewardson for performing lab work.

## Author Contributions

I.L. analyzed data with input from D.R. I.L., A.B-C., R.P., D.R. wrote the manuscript with input from other co-authors. A.B.-C., G.B.-O., O.B.-Y., N.J., E.K., D.L., Z.M. T.M. undertook archaeological work in Dzudzuana cave; G.B.-O. studied the archaeozoology of the cave and E.K. its palynology. S.M. performed bioinformatics. N.P. aided in the admixture graph analysis. O.C. and N.R. performed ancient DNA work. B.J.C. and D.J.K. performed ^14^C dating. R.P. and D.R. co-ordinated the work.

## Author Information

Correspondence and requests for materials should be addressed to I.L. or R.P., or D.R.

## Data and Code availability

The aligned sequences are available through the European Nucleotide Archive under accession number xxx. Genotype datasets used in analysis are available at https://reich.hms.harvard.edu/datasets. Code used to implement automated qpGraph fitting can be obtained from I.L. by request. All other data are available from the corresponding authors upon reasonable request.

